# Computational Prediction of Binding Affinities of Human Angiotensin Converting Enzyme-2 with SARS-CoV-2 Spike Protein Variants: Omicron Variants and Potentially Deleterious Mutations

**DOI:** 10.1101/2022.10.14.512203

**Authors:** Alexander H. Williams, Chang-Guo Zhan

## Abstract

The Omicron variant (BA.1) and its sub-variants of the SARS-CoV-2 virus which causes the COVID-19 disease continues to spread across the United States and the World at large. As new sub-variants of SARS-CoV-2 continue to proliferate, a reliable computational method of quickly determining the potential infectivity of these new variants is needed to assess their potential threat. In the present study, we have tested and validated an efficient computational protocol, which includes an efficient energy minimization and subsequent molecular mechanics/Poisson Boltzmann surface area (MM-PBSA) calculation of the binding free energy between the SARS-CoV-2 spike protein and human angiotensin converting enzyme-2 (ACE2), to predict the binding affinities of these spike/ACE2 complexes based upon the calculated binding free energies and a previously calibrated linear correlation relationship. The predicted binding affinities are in good agreement with available experimental data including those for Omicron variants, suggesting that the predictions based on this protocol should be reasonable. Further, we have investigated several hundred potential mutations of both the wildtype and Omicron variants of the SARS-CoV-2 spike protein. Based on the predicted binding affinity data, we have identified several mutations that have the potential to vastly increase the binding affinity of the spike protein to ACE2 within both the wildtype and Omicron variants.

**Author Summary:** As well known, the coronavirus responsible for COVID-19 disease enters human cells through its spike protein binding with a human receptor protein known as angiotensin converting enzyme-2. So, the binding affinity between the spike protein and angiotensin converting enzyme-2 contributes to the infectivity of the coronavirus and its variants. In this study, we demonstrated that a generally applicable, fast and easy-to-use computational protocol was able to accurately predict the binding affinity of angiotensin converting enzyme-2 with spike protein of the currently known variants of the coronavirus. Hence, we believe that this computational protocol may be used to reliably predict the binding affinity of angiotensin converting enzyme-2 with spike protein of new variants to be identified in the future. Using this computational protocol, we have further examined a number of possible single mutations on the spike protein of both the wildtype and Omicron variants and predicted their binding affinity with angiotensin converting enzyme-2, demonstrating that several mutations have the potential to vastly increase the binding affinity of the spike protein to angiotensin converting enzyme-2.

## Introduction

As the world enters the third year of the COVID-19 pandemic (caused by SARS-CoV-2), over five hundred million cases have been reported, with over six million deaths attributed to the disease. The virus’s ability to infect and destroy its host eclipses that of previous *Coronaviridae* such as SARS-CoV-1 (colloquially known as SARS), of which only infected 8,000 people across over 25 countries.[1] Pivotal to the infectivity of this virus is the spike protein (Gene S), which facilitates capsid attachment and integration into the host cell *via* attachment to membrane bound human angiotensin-converting enzyme II (ACE2), which is prolifically expressed within the respiratory system. The receptor binding domain (RBD) is the part of the spike protein which directly interacts with the ACE2 protein, containing a highly specialized receptor binding motif (RBM) which affords it a low-nanomolar binding affinity (22 nM) with ACE2. This binding affinity helps afford the SARS-CoV-2 virus its notable infectivity, and thus any potential changes to the RBD region of the spike protein could enhance this infectivity, which was seen with several variants of SARS-CoV-2 that appeared mere months after the initial spread of the virus.

As the pandemic continues to spread, many different strains of the virus have become dominant over the various regions of the globe, notable examples include the B.1.1.7 (Alpha variant) within the United Kingdom, B.1.617.2 (Delta variant) within the United States and the most recent Omicron (BA.1/BA.2) which caused a spike within United States in late 2021, resulting in a record 800,000 cases/day in mid-January 2022. All these mentioned variants contain modifications to both the RBD and RBM regions of the spike protein, resulting in changes to the binding affinity with the ACE2 protein.[2–9] The speed of which mutations have been incorporated into the spike protein has been blinding, with the first notable mutation, D614G, represented in over 25% of all sequenced samples before the virus even reached the United States.[10] However, determining which of these mutations poses a threat to the healthcare community would require thousands of variants to be synthesized and subsequently tested for their infectivity, a monumental stress on both time and resources. The new sub-strain of Omicron BA.4/5 presents a great threat, due to its new mutation L452R, a mutation previously seen within the Delta variant which confers significant evasiveness to many immune response antibodies as well as the clinically available antibodies such as Ly-CoV555.

Since the beginning of the pandemic, our lab has focused on the development of computational methodologies to predict the binding affinity of the Spike protein with various therapeutic targets or candidates, including the ACE2 protein, ACE2 mimics, as well as miniprotein inhibitors of the spike protein.[11, 12] Our efforts focused on utilizing accurate and computationally inexpensive methodologies such as energy minimization and molecular mechanics/Poisson Boltzmann surface area (MM-PBSA) calculations to estimate the binding free energy (ΔG) of these proteins/variants, and subsequently create a linear regression model that could be used on the variants with unknown binding affinities. As the Alpha variant began spreading throughout the United Kingdom,[13, 14] we applied this methodology to the spike protein, modifying the crystal structure of the wildtype spike/ACE2 complex that had been released, predicting a binding affinity of 0.44 nM. Before our results were even published, our prediction had been proven accurate during the review process, as the first measurement of the Alpha variant’s binding affinity was published at 0.8 nM.[11] As additional binding affinities and crystal structures for the SARS-CoV-2 variants were released, we continued to refine and use our linear regression model, predicting the binding affinity for the Omicron variant at 22.69 nM which was later validated at 20.6 nM.[15] With this improved model in-hand, the prediction of the binding affinities for mutated spike proteins with ACE2 became an inexpensive calculation, which could be performed in under 10 minutes per variant using a 32-Core processor.

As the first wave of the original Omicron variant (B.1.1.529 or BA.1) has waned, new offshoots of Omicron have taken hold in many different countries with BA.2 leading the charge. The Omicron variant represents a large change to the overall spike protein, with over 40 mutations applied *vs* the original spike protein encountered in the initial wave of the pandemic. These mutations have vastly increased the Omicron variant’s infectivity as well as provided evasiveness to immune responses and to clinically approved antibodies. Since the introduction of our generalized energy minimization/MM-PBSA methodology for predicting the binding affinity of these SARS-CoV-2 variants, empirical studies have been performed to obtain the dissociation constants of these variants’ spike proteins with human ACE2. This collection of binding affinity data presents an opportunity to further test (and refine, if necessary) our computational methodology for the prediction of binding affinities with the Omicron variant and its related substrains, which have overtaken the vast majority of reported SARS-CoV-2 cases.

To investigate both single point mutations of the spike RBD region as well as potential combinatorial mutations of the Omicron variant with other variants’ mutations (e.g., Beta, Delta, etc.), we employed our generalized energy minimization/MM-PBSA methodology to roughly 1,350 different variants of the spike protein, including the newly discovered Omicron sub-variants (BA.4/5 whose experimental affinity with ACE2 is known and BA.2.75 whose experimental binding affinity with ACE2 is unknown), and subsequently predicting their binding affinity using our previously established linear regression model.

## Methods

Our methodology utilizing energy minimization/MM-PBSA to determine the changes in binding affinity has been used with multiple types of therapeutics including small-molecule based inhibitors,[16–22] engineered-protein therapeutics,[23–25] and with multiple variants of concern (VOC) of the SARS-CoV-2 virus.[11, 26] This methodology has many advantages over other state-of-the-art techniques, the primary of which are the substantial savings in computational costs to predict the binding affinity of each pair of spike protein mutant and ACE2 protein. As opposed to an all-atom simulation[27–29] of the spike/ACE2 complex, which would require expensive CUDA-processing capable devices to perform multi-nanosecond (i.e., >10 ns) simulations to accurately determine the binding affinity, the energy minimization/MM-PBSA methodology can run on nearly any system with a capable x86 processor, with a prediction available for the binding affinity within 10 minutes for systems with 32-cores or more available. This methodology relies on the assumption that the crystal structure used within the calculations is close to its equilibrium or “unperturbed” state, and that the changes made to this system represent a small change in the overall structure, which can be returned to back to its “unperturbed” state *via* energy minimization. Additionally, our lab has previously shown that our “perturbation” method can accurately predict the binding affinity in both wildtype to mutated states and vice versa.[15]

To predict the binding affinity of the newly identified Omicron sub-variants, each variant’s sequence was obtained from the GISAID repository, and the S gene mutations were applied to the spike/ACE2 complex crystal structure (PDB: 7KMB)[30] using the PyMol mutagenesis tool to select the lowest energy rotamer of the mutated residues. Using pdb4amber, any non-protein entity was stripped of the PDB file, and the Amber topology and coordinates were prepared using the LEaP module of the AmberTools2020 package, using the ff99SB force field.[17, 18, 31–33] Each structure was then energy minimized using a two-step procedure, each consisting of 2,000 steps of steepest descent minimization and 3,000 steps of conjugate gradient descent, with each step placing 10 kcal/Å^3^ restraints on all non-hydrogen atoms and a 2 kcal/Å^3^ restraint on the backbone C_α_ atoms respectively. Finally, the binding free energy (ΔG) was estimated using the MMPBSA.py module of the AmberTools package, using the built in quasi-harmonic methodology to estimate the entropy for the binding mode.

To predict the binding affinity of hundreds of single residue mutations upon the spike protein, the crystal structure of the spike/ACE2 complex (PDB: 7KMB)[30] was loaded into PyMol and the entire ACE2 protein was selected, this selection was expanded to include all residues upon the spike protein within 6 angstroms of ACE2. These residues were then mutated to each standard amino acid using the FoldX BuildModel module, choosing the lowest energy rotamer for each mutation Using pdb4amber, any non-protein entity was stripped of the PDB file, and the Amber topology and coordinates were prepared using the LEaP module of the AmberTools2020 package, using the ff99SB force field. Each structure was then energy-minimized using a two-step procedure, each consisting of 2,000 steps of steepest descent minimization and 3,000 steps of conjugate gradient descent, with each step placing 10 kcal/Å^3^ restraints on all non-hydrogen atoms and a 2 kcal/Å^3^ restraint on the backbone C_α_ atoms respectively. Then, the binding free energy was estimated using the MMPBSA.py module of the AmberTools package, using the built in quasi-harmonic methodology to estimate the entropy for the binding mode. The predicted binding affinity was then corrected using our previously published linear regression model to account for the typical overestimation seen within the MM-PBSA methodology.[11, 15] Similarly, the dual residue mutations were created by identifying residue changes within other variants of concern (VOCs), that are not present within the Omicron variant (e.g., L452R and T478K from the Delta variant, N501Y from the Alpha variant, etc.), and creating each combination of two VOC mutations and applying them to the Omicron variant structure using the FoldX BuildModel module, which were then run through the same energy minimization/MM-PBSA methodology as the single residue mutation structures.

Finally, the binding free energies (Δ*G*_PB_) obtained from the MM-PBSA calculation were corrected by using the previously reported empirical equation (Eq 1).[15] The corrected binding free energy (Δ*G*_corr_) is our prediction for binding affinity of the spike variant with ACE2.

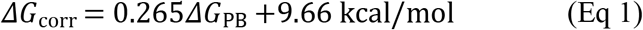

## Results

### Omicron (BA) lineages of SARS-CoV-2

The binding free energies calculated for the Omicron variants and previously known variants are summarized in Table 1, in comparison with available experimental data. As seen in Table 1, the predicted binding affinities using our previously published empirical equation (Eq. 1)[15] are reasonably close to the corresponding experimental data, with a root-mean-square error (RMSE) of 0.38 kcal/mol. The reasonable agreement between the predicted binding affinities and the experimental data further validated our computational protocol, suggesting that the predicted binding affinity (*K*_d_ = 19.47 nM or ~20 nM based on Eq 1) for the BA.2.75 variant (whose experimental binding affinity has not been reported at this point) should be reasonable.

**Table 1.**
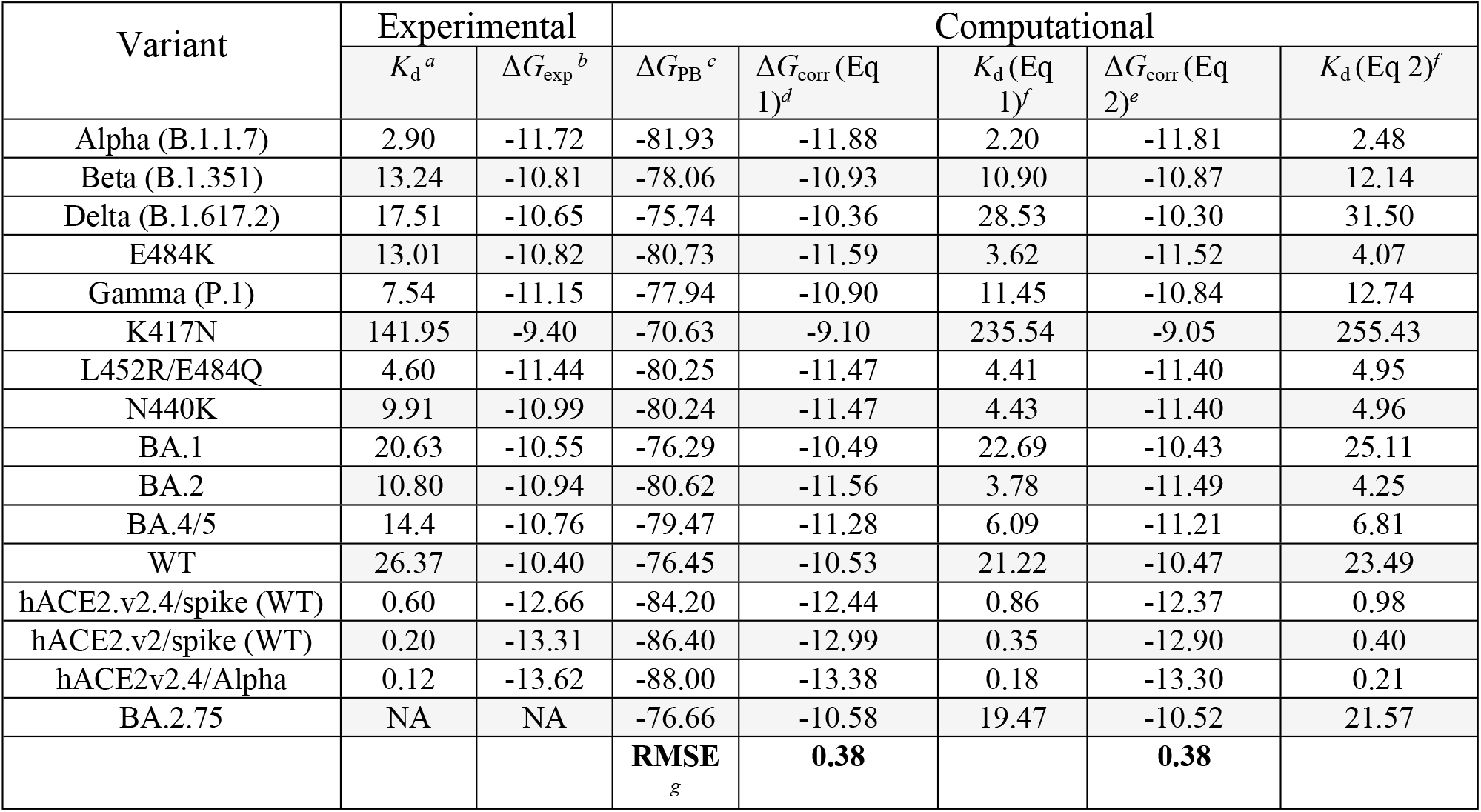
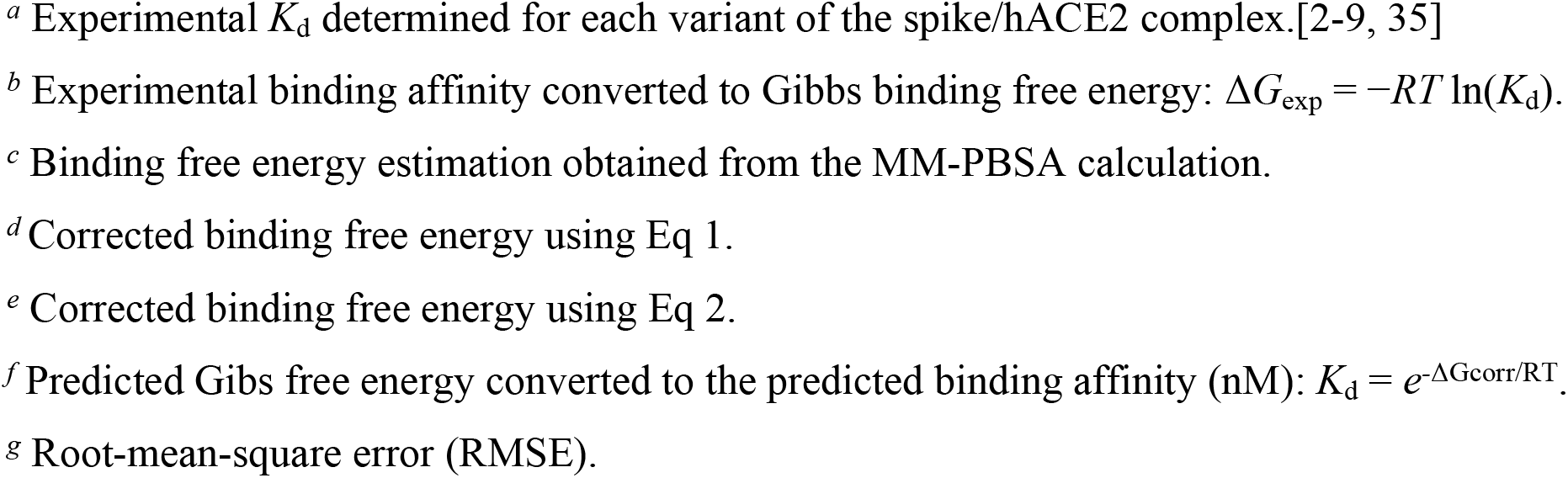
Comparison of the predicted binding free energy (kcal/mol) and *K*_d_ (nM) values *vs* the experimentally determined binding affinity.

| Variant | Experimental |  | Computational |  |  |  |  |
| --- | --- | --- | --- | --- | --- | --- | --- |
| | $K_d^a$ | $\Delta G_{\text{exp}}^b$ | $\Delta G_{\text{PB}}^c$ | $\Delta G_{\text{corr}}(\text{Eq } 1)^d$ | $K_d(\text{Eq } 1)^f$ | $\Delta G_{\text{corr}}(\text{Eq } 2)^e$ | $K_d(\text{Eq } 2)^f$ |
| Alpha (B.1.1.7) | 2.90 | -11.72 | -81.93 | -11.88 | 2.20 | -11.81 | 2.48 |
| Beta (B.1.351) | 13.24 | -10.81 | -78.06 | -10.93 | 10.90 | -10.87 | 12.14 |
| Delta (B.1.617.2) | 17.51 | -10.65 | -75.74 | -10.36 | 28.53 | -10.30 | 31.50 |
| E484K | 13.01 | -10.82 | -80.73 | -11.59 | 3.62 | -11.52 | 4.07 |
| Gamma (P.1) | 7.54 | -11.15 | -77.94 | -10.90 | 11.45 | -10.84 | 12.74 |
| K417N | 141.95 | -9.40 | -70.63 | -9.10 | 235.54 | -9.05 | 255.43 |
| L452R/E484Q | 4.60 | -11.44 | -80.25 | -11.47 | 4.41 | -11.40 | 4.95 |
| N440K | 9.91 | -10.99 | -80.24 | -11.47 | 4.43 | -11.40 | 4.96 |
| BA.1 | 20.63 | -10.55 | -76.29 | -10.49 | 22.69 | -10.43 | 25.11 |
| BA.2 | 10.80 | -10.94 | -80.62 | -11.56 | 3.78 | -11.49 | 4.25 |
| BA.4/5 | 14.4 | -10.76 | -79.47 | -11.28 | 6.09 | -11.21 | 6.81 |
| WT | 26.37 | -10.40 | -76.45 | -10.53 | 21.22 | -10.47 | 23.49 |
| hACE2.v2.4/spike (WT) | 0.60 | -12.66 | -84.20 | -12.44 | 0.86 | -12.37 | 0.98 |
| hACE2.v2/spike (WT) | 0.20 | -13.31 | -86.40 | -12.99 | 0.35 | -12.90 | 0.40 |
| hACE2v2.4/Alpha | 0.12 | -13.62 | -88.00 | -13.38 | 0.18 | -13.30 | 0.21 |
| BA.2.75 | NA | NA | -76.66 | -10.58 | 19.47 | -10.52 | 21.57 |
|  |  |  | **RMSE<sub>g</sub>** | **0.38** |  | **0.38** |  |

- <sup>a</sup> Experimental $K_d$ determined for each variant of the spike/hACE2 complex.[2-9, 35] - <sup>b</sup> Experimental binding affinity converted to Gibbs binding free energy: $\Delta G_{\text{exp}} = -RT \ln(K_d)$ . - <sup>c</sup> Binding free energy estimation obtained from the MM-PBSA calculation. - <sup>d</sup> Corrected binding free energy using Eq 1. - <sup>e</sup> Corrected binding free energy using Eq 2. - <sup>f</sup> Predicted Gibbs free energy converted to the predicted binding affinity (nM): $K_d = e^{-\Delta G_{\text{corr}}/RT}$ . - <sup>g</sup> Root-mean-square error (RMSE).

BA.2.75 variant[34] is a subvariant of BA.2 with several additional mutations including D339H, G446S, N460K, and R493Q, the last of which is in close proximity to the binding interface with ACE2 (Figure 1). Using our mutation/energy minimization/MM-PBSA methodology along with Eq 1, we have found that the mutations seen within this new variant impose a disadvantage in binding to the ACE2 protein over its parent BA.2 subvariant of Omicron. This decrease in binding affinity primarily comes from the mutation of R493Q which replaces a positively charged arginine that is within close proximity to two negatively charged glutamic acid residues (E35 and E38) within ACE2. Replacing this residue with glutamine removes a strong synergistic charge/charge interaction with the negatively charged E35 (Figure 1). However, this loss in a strong charge interaction is partially mitigated by the hydrogen bond that is formed between the glutamine residue’s sidechain and the backbone oxygen of E35 (Figure 1). Further experimental study will be required to validate this prediction in binding affinity for BA.2.75.

**Figure 1.**
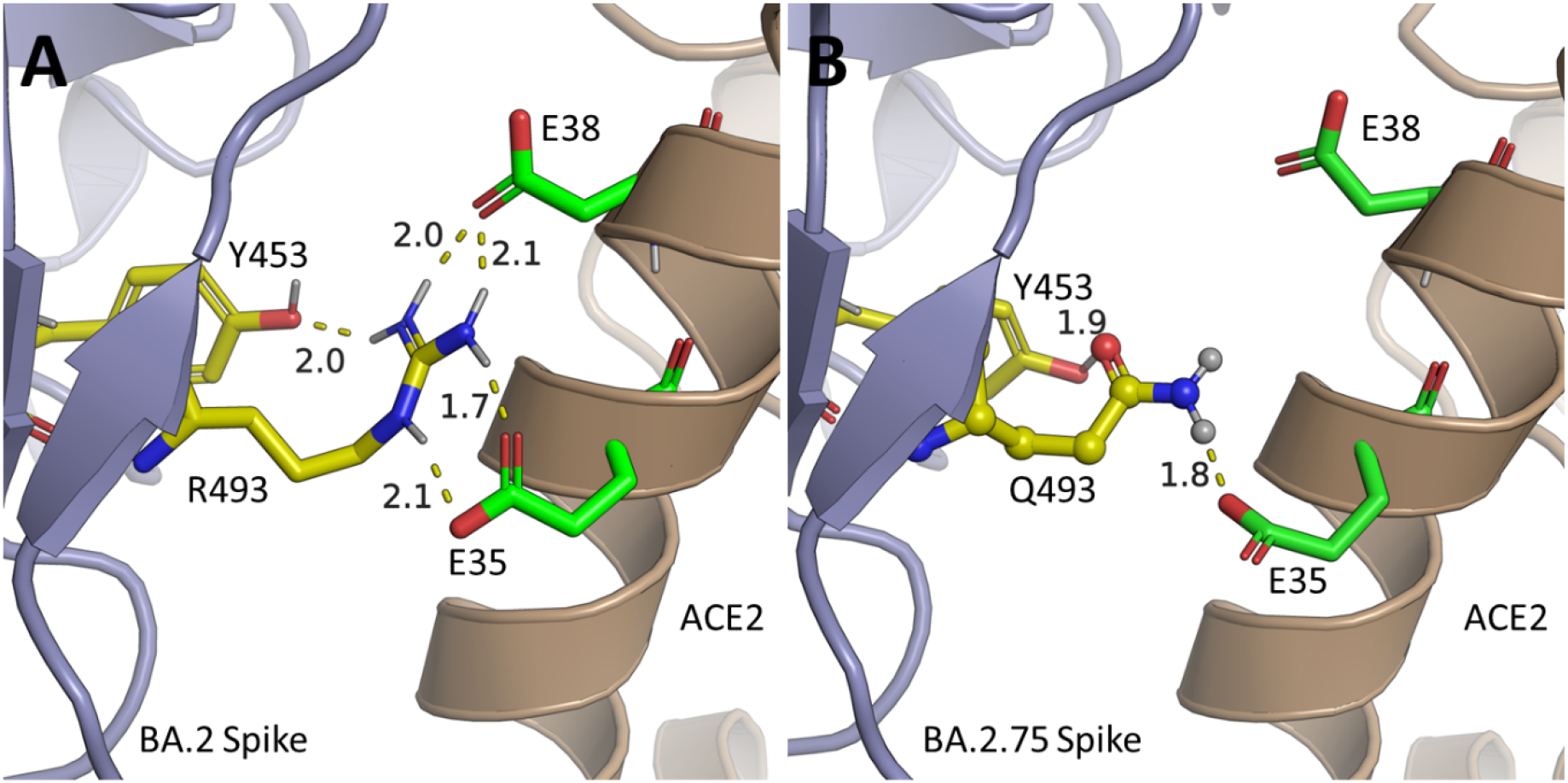
Binding mode of the BA.2 (**A**) and BA.2.75 (**B**) variants of the SARS-CoV-2 spike protein with human ACE2. The Q493R mutation found within BA.2 is reversed within BA.2.75 to the original Omicron Q493 (represented in ball and stick), this mutation removes a strong charge interaction with nearby negatively charged residue E35 and E38, leading to the predicted loss in binding affinity between the BA.2.75 variant and ACE2. However, a strong hydrogen bond can form between the Q493 sidechain and the sidechain of E35, mitigating a total loss in interactions between the two residues.

Further, with more experimental binding affinity data (summarized Table 1) available now, we have also recalibrated the linear regression equation and obtained the new model (Eq 2):

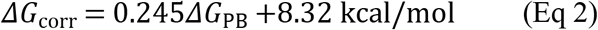

Depicted in Figure 2 are the plots of the experimental binding free energy *vs* the predicted binding free energy using Eq 1 or Eq 2. It turns out that both Eq 1 and Eq 2 are similarly accurate. For the overall accuracy, both equations have the same correlation coefficient (*R*^2^ = 0.894) and RMSE (0.38 kcal/mol), as shown in Figure 2. For this reason, all other predictions to be discussed below were based on the use of Eq 1.

**Figure 2.**
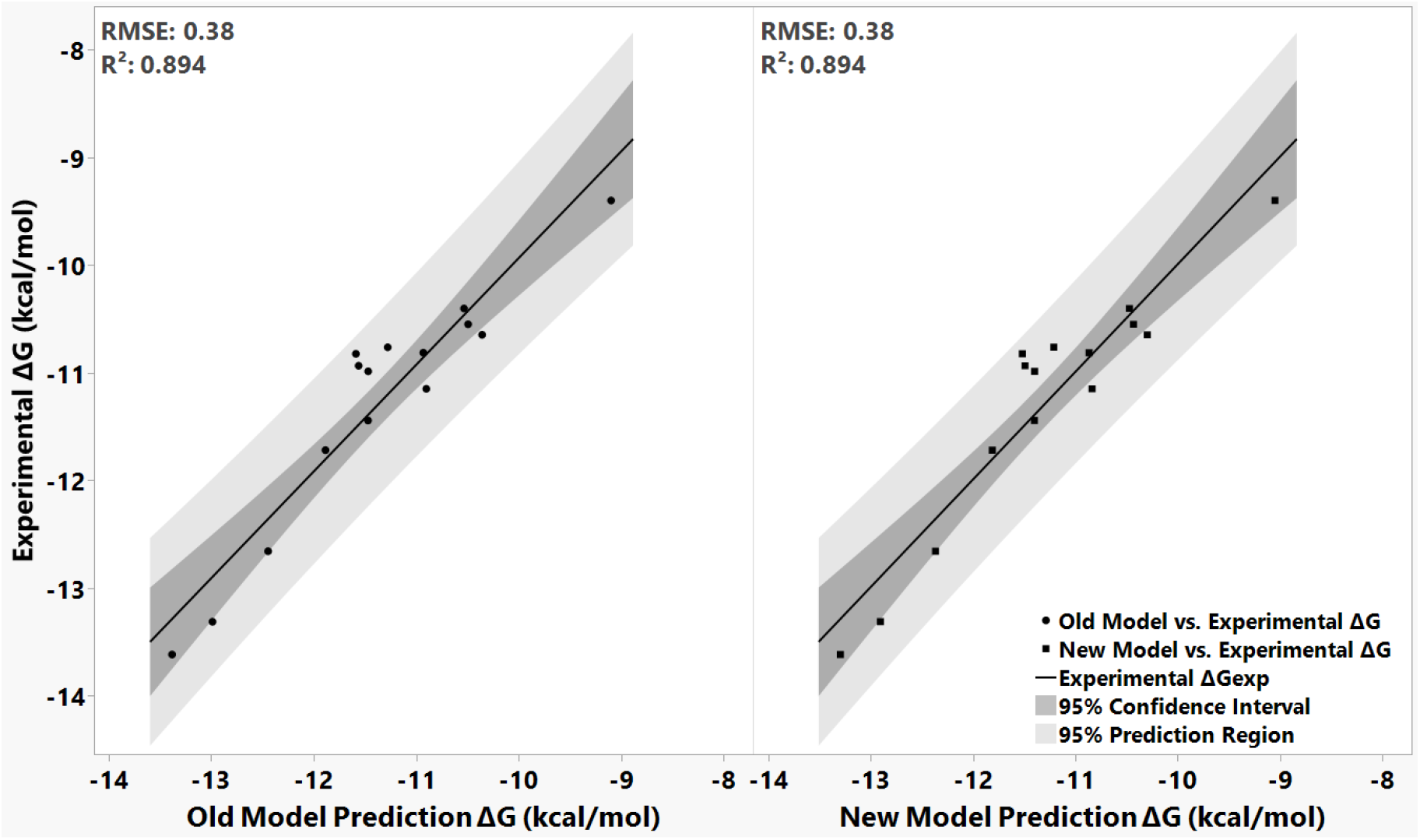
Plots of the experimental binding free energy (ΔG_exp_) vs the predicted binding free energy (ΔG_corr_) corrected by using Eq 1 (old model prediction) or Eq 2 (new model prediction). The dark shaded region represents the 95% confidence interval of the predictions, whilst the light shaded region represents the 95% prediction region of where all predictions occur.

### *Single-Point Mutations of the* W*ildtype Spike Protein*

By choosing spike residues within six angstroms of the ACE2 protein, 39 residues in proximity were selected for modification. By mutating these residues to every standard amino acid, a total of 780 mutated spike proteins were generated. The predicted binding affinity of the spike variants with the single residue mutations can be found in Figure 3, with additional details available in Table S1 (Supporting Information). An in-depth representation of the binding mode of several notable residue mutations can be found in Figure 4.

According to predicted binding affinity data shown in Figure 3, of these 780 mutations, 361 had a predicted stronger binding affinity with ACE2 when compared to the wildtype protein, 104 had a similar binding affinity (within the RMSD of the linear regression model[15]), and 315 mutations had a weaker binding affinity to ACE2. Of the positions tested, R403 was the most precarious, with most residue changes resulting in a large positive change in the ΔG predicted by the MM-PBSA methodology, which comes from the loss in internal stability within the spike protein (Figure 1).

**Figure 1.**
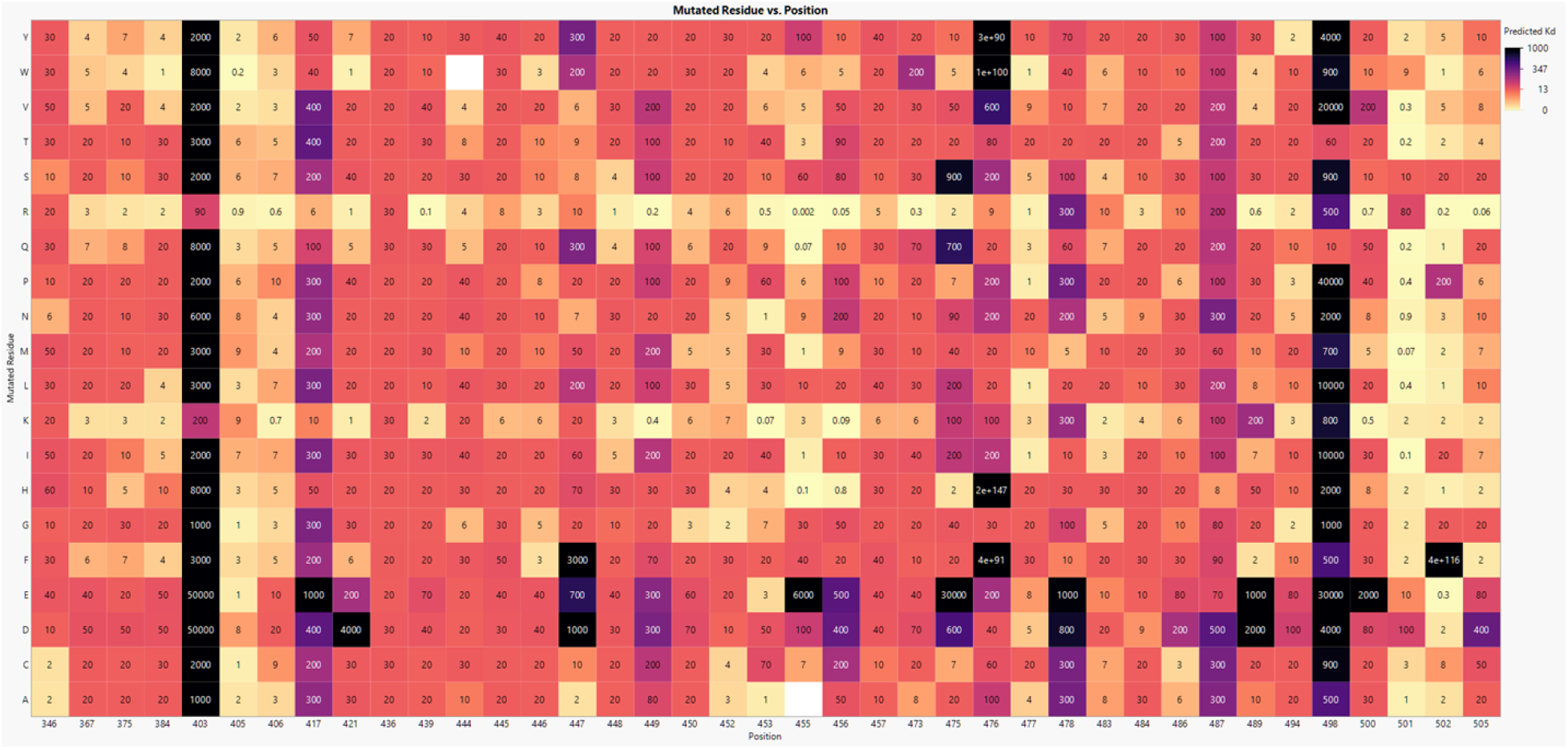
Predicted binding affinity *K*_d_ (in nM) of 780 mutations of the wildtype spike protein. Darker colors indicate weaker binding affinities. Mutations to R403 display the worst affinity to the ACE2 protein due to contributing internal steric strain to the spike protein. Only the flexible and positively charged lysine and arginine residues can fit within this space.

This large change can primarily be attributed to the enclosed area surrounding R403, which requires the flexibility of the long arginine residue to avoid steric clashing with other residues of the spike protein (Figure 2A). Additionally, negatively charged residues such as aspartic and glutamic acid completely change the charge of the residue, causing negative charge repulsion with several residues in the periphery of R403, including D405 and E37. Other residues showed more affinity to being mutated, including L452 and L455 both of which showed considerable increases in binding affinity when mutated to positively charged residues arginine and histidine; L452R is a common mutation and is most prolifically seen within the B.1.617.2 or Delta variant; the mutation to the positively charged arginine residue compliments the overall negatively charged ACE2 protein’s E35, E37, and D38. The mutation of L455R similarly introduces +/- charge interactions with several negatively charged residues including D30 and E35 (Figure 2B), and the opposite effect can be observed when mutating these residues to negatively charged residues such as aspartic and glutamic acid, which decreases their binding affinity significantly (100 and 6000 nM, respectively.)

**Figure 2.**
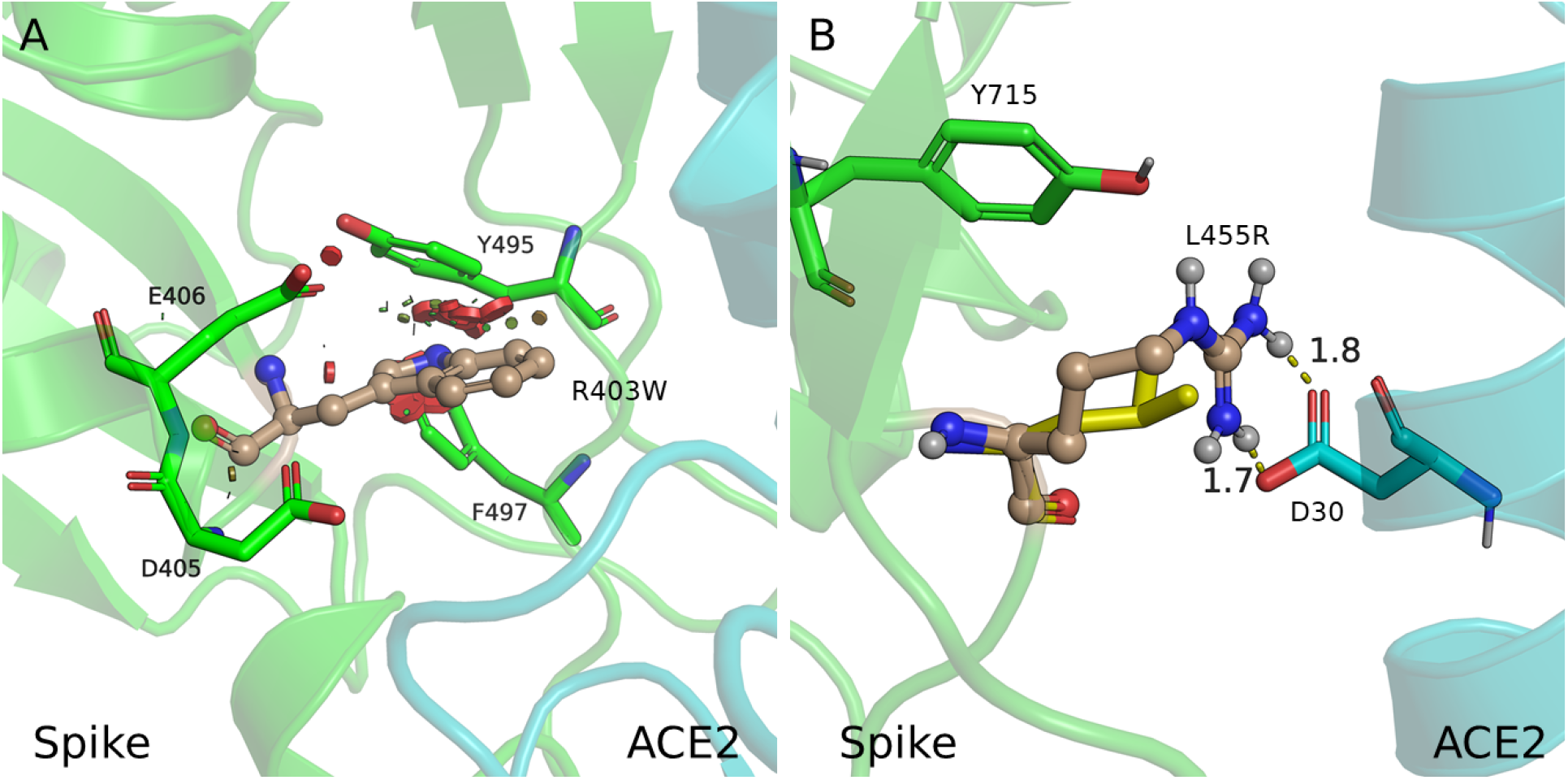
(A) Binding mode of the R403W mutation (predicted *K*_d_ = 8000 nM) with nearby residues highlighted in stick representation. Tryptophan is in its lowest energy rotamer within this space, and sterically clashes with Y495 of the spike protein. Areas of steric strain are highlighted with red discs. (B) The L455R mutation, which had the strongest predicted binding affinity of all tested wildtype mutations. This high affinity results from the strong +/- charge interactions created with nearby residue D30, replacing the mismatched interactions between the hydrophobic L455 on the spike protein and hydrophilic D30 upon ACE2.

### Combinatorial Mutations of the Omicron Variant Spike Protein

The healthcare community has begun taking note of several combinatorial variants that have been discovered within France and the United Kingdom. While these variants (e.g., XD, XE, and XF) do not have crossover within their spike protein gene, the potential for infective variants such as Omicron gaining the mutations of more deadly variants such as Delta’s deserve further investigation to see if their mutations are synergistic. Hence, we have also examined the other possible dual residue mutations mentioned above (in the Introduction section). By utilizing Eq 1, we have predicted the binding affinities for a variety of dual residue mutations, and the predicted data can be found in Figure 5 with additional details available in Table S2. Based on the binding affinity predictions, we have identified several dual mutations that we should be vigilant for. The most prominent changes upon the spike protein for the Delta variant are the L452R and T478K mutation, both located upon the RBM of the spike protein. The L452R mutation is notably absent upon the Omicron variant’s spike protein and as such could present a way for its spike protein to gain considerable binding affinity with ACE2. The L452R mutation by itself (shown as the diagonal in Figure 5, in which double mutations consist of the same mutation twice), shows the potential to increase the binding affinity of the Omicron variant’s spike protein to 4.6 nM, a roughly 5-fold increase in binding affinity.

Other notable examples include the double mutations which include the N439K mutation, which mimics the same interactions seen within the N440K mutation also contained within the Omicron variant. These two mutations can create a +/- charge interaction with the nearby E329 residue upon ACE2; however, N339K can increase the strength of this interaction due to being closer in proximity to the ACE2 protein. Additionally, N339K can donate a hydrogen bond with nearby Q325 in ACE2 further strengthening the two proteins’ interactions. These two strong interactions allow for the N339K mutation to obtain some of the strongest binding affinities of all the double mutations studied as this mutation by itself increases the binding affinity to 1.4 nM overall. The top double mutation also contains the E484A mutation which rectifies another mutation within the Omicron variant; the Q493R mutation of the Omicron variant introduces both strong hydrogen bond and charge interactions with E35 of ACE2, however this mutation also weakens an intra-protein interaction between E35 and K31 within ACE2, as the positive charge of the mutated arginine Q493R repels the also positively charged lysine residue. This movement additionally weakens the interactions between E484 and K31 between the spike and ACE2 proteins (Figure 4B); the mutation to E484A removes this now unpaired glutamic acid and removes the repelling force of the overall negatively charged ACE2 protein.

**Figure 3.**
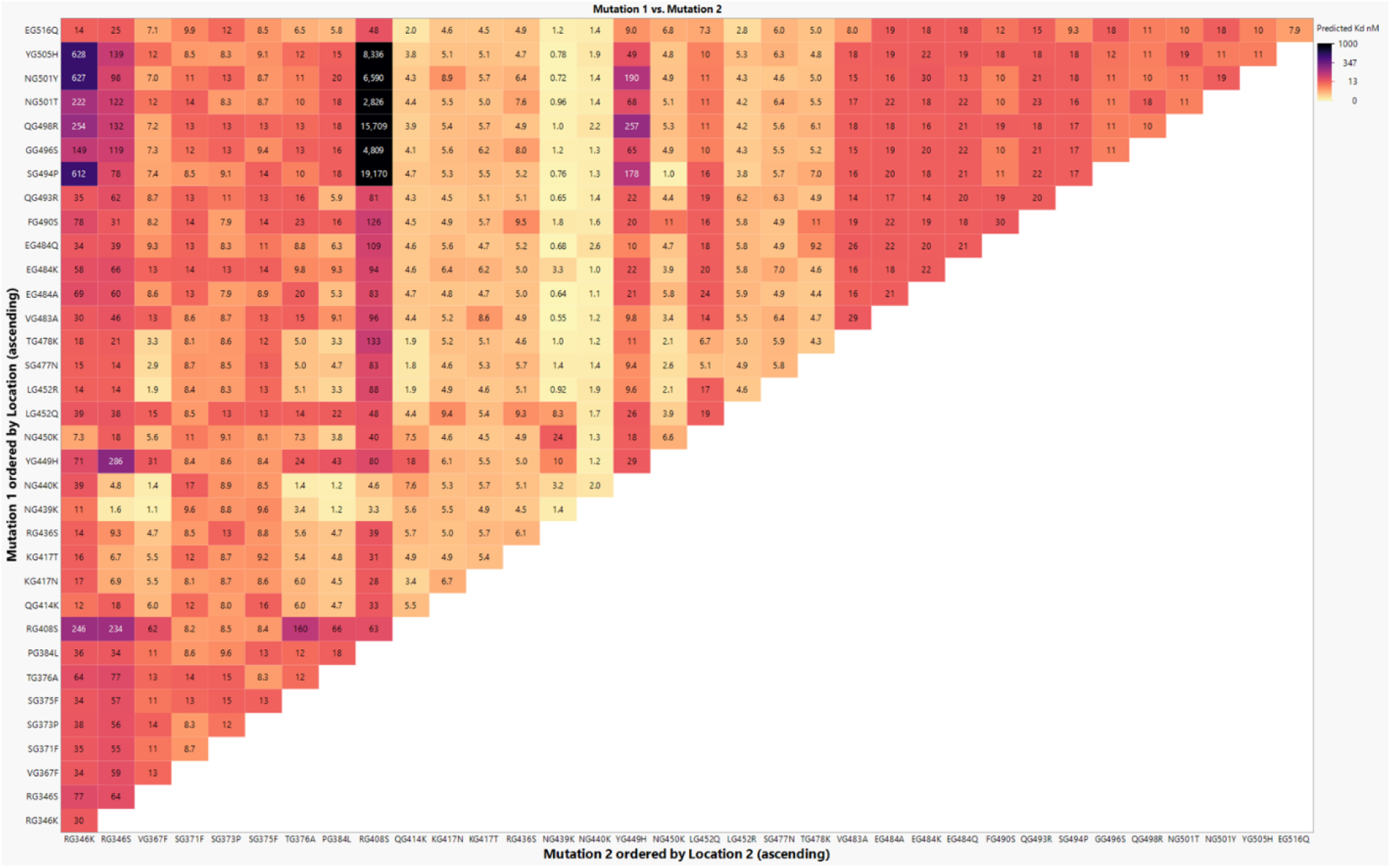
Predicted binding affinities of all double mutations upon the Omicron variant, with darker colors representing weaker binding affinity with the ACE2 protein. Double mutations along the diagonal contain the same mutation, resulting in a single mutation upon the Omicron variant. N439K and N440K represent the mutations with the strongest average binding affinity due to their creation of a strong +/- charge interaction with ACE2.

**Figure 4.**
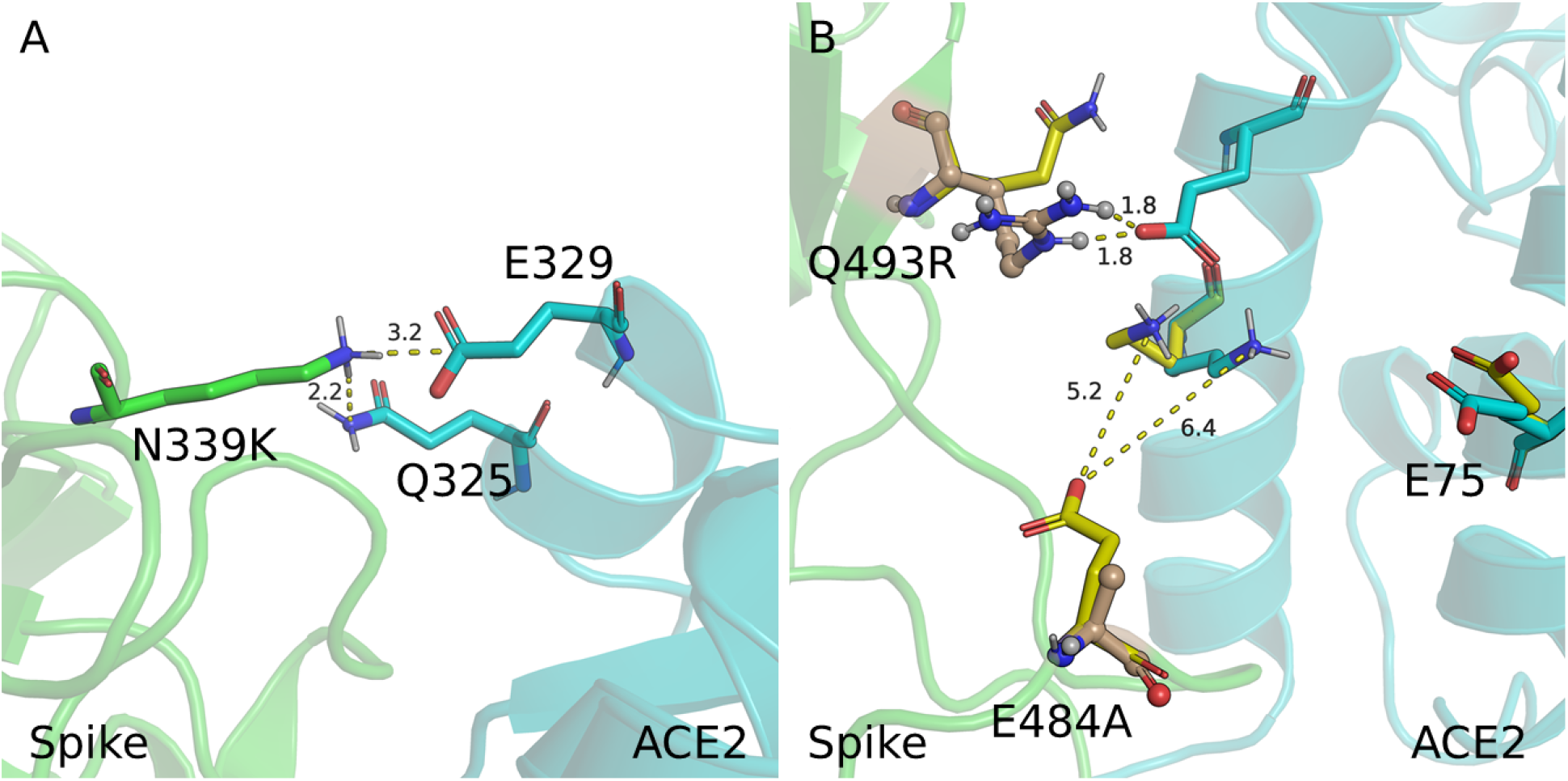
(A) Contribution of the N339K mutation to the Binding mode of the Omicron variant, creating a strong hydrogen bond with Q325 upon ACE2 and a +/- charge interaction with E329; these creation of these two interactions allows the N339K mutation alone to increase the binding affinity of the Omicron variant to ACE2 14-fold. (B) Contribution of the E484A mutation to the N439K/E484A double mutant of the Omicron variant. The shift of glutamic acid to alanine replaces an otherwise weakened interaction of E484 to K31 upon ACE2, which increases the overall repulsive interactions between the negatively charged ACE2 and the negatively charged E484.

## Discussion

Through the use of our previously established, computationally efficient methodology for estimating the binding affinity of spike/ACE2 complexes using the energy minimization and MM-PBSA estimation of the binding free energy, we have tested and validated our model’s predictions to the Omicron variant’s sub-strains, including the ascendant BA.4/5 variant. Our methodology was able to produce binding free energy estimations that are reasonably close to the experimentally determined binding free energy values obtained for each sub-strain’s spike/ACE2 complex binding. Incorporating these additional empirically backed values into our linear regression model did not impact the original model’s (i.e., Eq.1) prediction correlation with the empirical data (R^2^) or the root-mean-square error value (RMSE) for all predictions. Using this same methodology, we have examined the newly reported BA.2.75 variant, and have predicted a lower binding affinity with ACE2 *vs*. its parent BA.2 strain, however this affinity is still on par with the original Omicron variant and should not be underestimated.

Additionally, we have predicted the binding affinity of over 1350 combined potential single and double mutations to the wildtype and Omicron variant spike protein. Overall, the negative charge of the ACE2 protein allows for mutations which introduce additional positive charges to the spike protein to increase the binding affinity of the complex, however this can be limited by steric strain introduced by the bulkiness of said charged residues (e.g., arginine residues). While the RBM region of the spike protein is an ideal location to introduce these positively charged residues, its proximity to the ACE2 excludes some residues from being mutated at all without severely affecting the binding affinity (e.g., Q498, Figure 3).

These predictions for the mutations of the Omicron spike protein identified L452R and N339K as potential mutations that could increase the binding affinity of the spike protein with ACE2. While the L452R mutation has been reported in the BA.4/5 sub-strain of the Omicron variant, additional mutations convolute our comparison with the experimental data for this single-point mutation of the Omicron variant. As the SARS-CoV-2 virus continues to mutate and spread, this study may be used as a reference to understand how each mutation will affect the binding affinity of the spike protein to ACE2 and by partial extension, its overall infectivity.

## Supporting Information

Additional tables, including Tables S1 (summarizing single mutation data) and S2 (summarizing double mutation data to the Omicron variant).

## Notes

The authors declare no competing financial interest.

## Acknowledgments

This work was supported in part by the funding of the Molecular Modeling and Biopharmaceutical Center at the University of Kentucky College of Pharmacy, the National Science Foundation (NSF grant CHE-1111761), and the National Institutes of Health (R01 DA056646-01). The authors also acknowledge the Computer Center at University of Kentucky for supercomputing time on their Lipscomb Compute Cluster, and their NVIDIA V100 nodes.

## Notes

### Competing Interest Statement

The authors have declared no competing interest.

